# Improved deconvolution of circulating tumor DNA from ultra-low-pass whole-genome methylation sequencing using CelFiE-ISH

**DOI:** 10.64898/2026.04.15.718632

**Authors:** Efrat Katsman, Sara Isaac, Alaa Darwish, Myriam Maoz, Michal Inbar, MazalTov Marouani, Irene Unterman, Ahinoam Gugenheim, Nada Salaymeh, Shatha Abu Khdeir, Beatrice Uziely, Tamar Peretz, Luna Kaduri, Ayala Hubert, Jonathan E Cohen, Azzam Salah, Mark Temper, Tamar Sela, Albert Grinshpun, Aviad Zick, Benjamin P. Berman, Amir Eden

## Abstract

Liquid biopsy using ultra-low-pass whole-genome sequencing (ULP-WGS, ∼0.25x coverage) is a promising tool to detect circulating tumor DNA (ctDNA) for cancer management, and the use of the native Oxford Nanopore (ONT) sequencing platform adds DNA methylation to the set of detectable features. Here, we test the performance of methylation-based cell-type deconvolution in ULP-WGS samples from diverse epithelial malignancies and investigate several new computational strategies using our CelFiE-ISH deconvolution framework. We find that incorporating larger numbers of markers restricted to the epithelial cell lineage can reduce the cancer fraction limit of detection down to 1.7-3.1%, matching or exceeding the 3% floor of established copy-number alteration (CNA) benchmarks. Our study provides a useful strategy for analysis of ULP-WGS ONT data and indicates that marker selection remains a key challenge for analyzing methylation-based cancer datasets.

## Introduction

DNA methylation is a highly informative cancer biomarker, but both tissue-based and blood-based clinical samples consist of complex cell type mixtures. A primary challenge in analyzing these data is the accurate decomposition, or deconvolution, of the constituent cell types. In oncology, methylation is used to identify circulating tumor DNA (ctDNA) from blood samples, and can reveal the type and molecular properties of cancer cells (1, 2). While methylation-based detection of ctDNA is widely utilized, it remains technically challenging because the cancer-derived fraction of total cell-free DNA (cfDNA) is often minute, often 5% or less at the time of diagnosis.

Reference-based deconvolution algorithms typically rely on a comprehensive atlas of DNA methylation signatures derived from a broad range of healthy cell types (3). A variety of computational frameworks have been developed to utilize these atlases, including the Expectation-Maximization-based CelFiE (4) and its fragment-level extensions CelFEER (5) and CelFiE-ISH (6), along with another fragment-level model, UXM (3). Deep-learning-based approaches such as cfSort (7) and MEnet (8) have also had encouraging results. These diverse methodologies have been subjected to rigorous systematic benchmarking (9), providing insights into their comparative performance across different sequencing depths and clinical contexts.

The selection of informative genomic markers is a critical determinant of deconvolution accuracy, as the choice of regions profoundly impacts model performance. While short, unmethylated regions are characteristic of cell-type-specific gene regulatory regions and enhancers (3), their specificities range from individual cell types (e.g., basal lung epithelial vs. lung pneumocyte epithelial) to broader cell type lineages such as pan-epithelial or pan-hematopoietic, to other combinations that can span multiple distinct lineages (7). Marker properties can be tailored to the algorithm; for haplotype-aware methods like CelFiE-ISH, high CpG density and "bimodal-ness"—quantified by the Bayesian Information Criterion (BIC)—are the strongest predictors of performance (6). Criteria further diverge between high-depth targeted sequencing (10) and low-coverage whole-genome sequencing (WGS). At ultra-low-pass WGS depths of 0.1–0.5x (ULP-WGS, (11)), deconvolution is challenging for rare tumor fractions (<1–5%) because many informative markers lack even a single tumor-derived read. While CelFiE-ISH can detect specific cell types down to 0.03% at 5x coverage, performance for all models declines markedly below this 5x threshold (6).

Oxford Nanopore Technologies (ONT) offers a rapid alternative for cfDNA analysis by natively profiling DNA methylation without chemical conversion, preserving DNA integrity and simplifying workflows for near-patient applications. Our recent work demonstrated that ultra-low-pass whole-genome ONT sequencing (∼0.2x–0.5x) could detect cancer-specific methylation, copy-number, and fragmentation signatures of ctDNA in plasma from metastatic lung cancer patients (12); however, most samples in that study had relatively high tumor fractions. Here, we perform low-coverage (0.2–0.3x) ONT sequencing on plasma samples from a range of epithelial malignancies, and across a range of tumor fractions, to investigate the impact of marker selection and improved deconvolution using CelFiE-ISH algorithm. We find that marker selection profoundly impacts the sensitivity and accuracy of ctDNA detection and that a method utilizing pan-epithelial markers provides superior detection in these low-coverage, low-tumor-fraction contexts, a finding that held for several other popular deconvolution algorithms we tested.

## Results

### CelFiE-ISH deconvolution is improved by clipping low-confidence cell type assignments

We extracted cell-free DNA from the plasma of 20 samples from five advanced colon or rectal adenocarcinoma patients and 22 non-cancer control donors and performed Oxford Nanopore (ONT) whole-genome sequencing to a median depth of 3.2M fragments (approx 0.22x genome coverage), using methods adapted from (12). As a baseline, we estimated tumor fraction in each sample from somatic copy number alterations (CNAs) using ichorCNA (11). ichorCNA was run using a panel of normals that included all healthy control samples plus additional “healthy-like” breast cancer control samples sequenced using the same protocol (Supplemental Figure 1A-B and Supplemental Table 1). The colon cancer samples had a median ichorCNA tumor fraction of 0.35.

We performed cell type deconvolution as previously described, using the CelFiE-ISH package and its reference atlas of 31 healthy cell types (6). For each cell type, we used the 250 unmethylated marker regions described in (3), excluding markers located on sex chromosomes, a total of 7451 marker regions across all cell types. We applied the three deconvolution methods implemented in this package, UXM, CelFiE, and CelFiE-ISH, as well as an additional method CelFEER (5). All four methods identified the expected cell types - monocytes, lymphocytes, erythroid/megakaryocyte (EP/MK), granulocytes (neutrophils), and small fractions of hepatocyte and/or endothelial cells (Figure 1A and Supplemental Figure 2).

**Figure 1.**
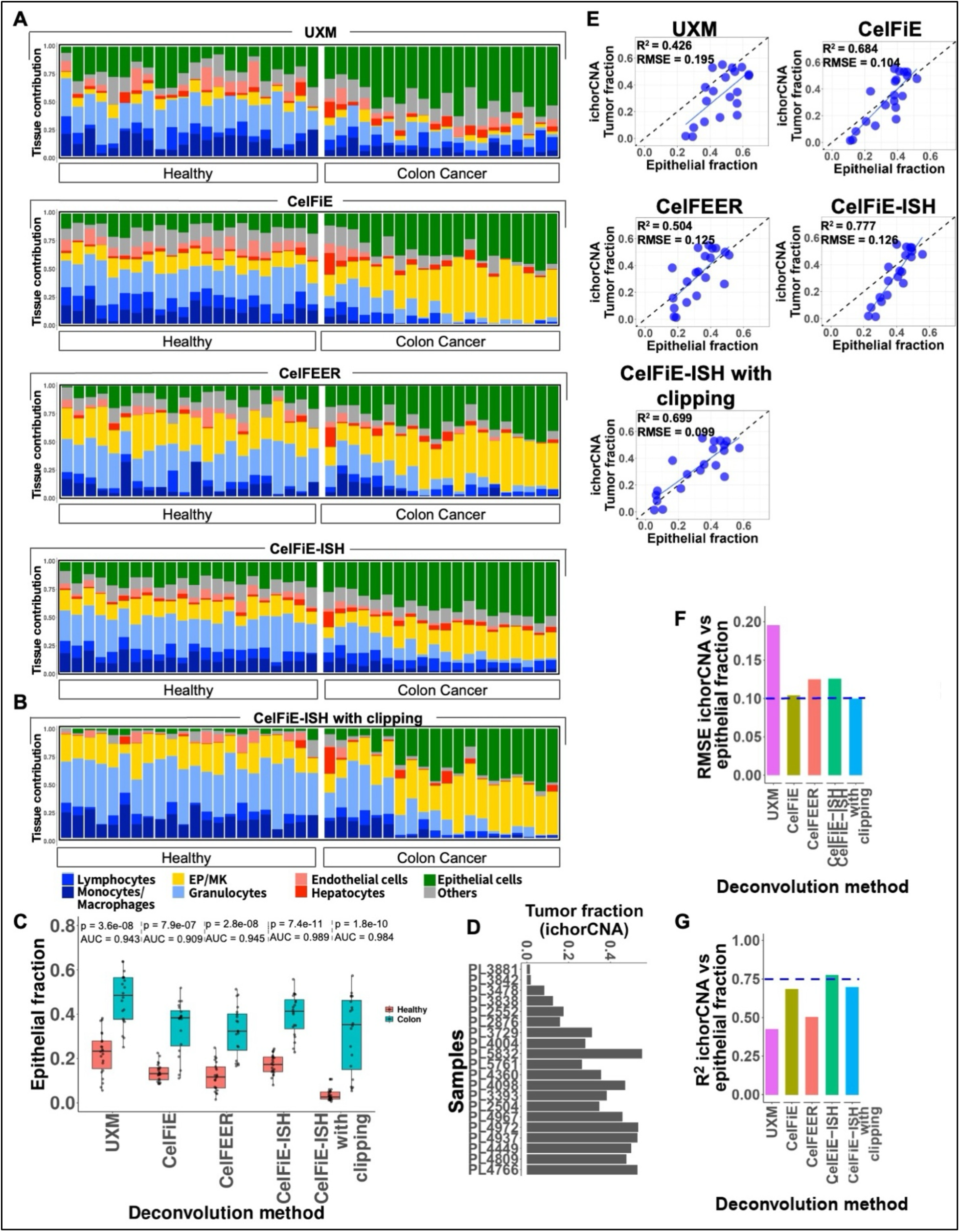
CelFiE-ISH deconvolution is improved by clipping low-confidence cell type assignments. **(A)** Cell-of-origin contributions were estimated using four computational deconvolution methods: UXM, CelFiE, CelFEER, and CelFiE-ISH, with 31 cell types and 250 markers per cell type as described in the text. Analyses were performed on healthy (n = 22) and colorectal cancer (n = 20) cohorts. Stacked bar plots display the proportional contributions of 31 cell types, and grouped into seven categories plus “Others” (sample identifiers are shown in Supplemental Figure 2). **(B)** Same visualization using CelFiE-ISH with clipping mode. **(C)** Epithelial fractions were compared between healthy and cancer samples separately for each deconvolution method, calculating AUROC and a statistical p-value using a one-tailed Wilcoxon test. **(D)** ichorCNA tumor fractions for the cancer samples, with the same sample ordering as above. **(E)** Comparison of methylation-based epithelial fractions against ichorCNA tumor fractions. **(F)** Root Mean Squared Error (RMSE) and **(G)** Correlation (R^2^) across the deconvolution methods.

We summed the fractions of all epithelial cell types into a single composite “epithelial” cell type, as in (12) (Figure 1A and Supplemental Figure 2). While the epithelial fraction was high in the cancer samples as expected, it was also higher than expected in the non-cancer controls, which should have very little epithelial content (3). Inflated estimates for non-existent cell types in these deconvolution tools were noted in the CelFiE-ISH paper (6), which speculated that they may be the result of low-confidence cell-type assignments for individual reads. To test this, we took advantage of the probabilistic behavior of the CelFiE-ISH deconvolution, by setting any low-confidence cell-type assignment (those with probabilities <0.05) to 0. The modified CelFiE-ISH version (hereafter “CelFiE-ISH with clipping”) reduced the epithelial component in the control samples to near-zero levels, while leaving most of the cancer samples with a significant epithelial component (Figure 1B). The version with clipping also reduced the contribution of the cell types labeled “Others”, which are not typically found in healthy plasma (shown in detail in Supplemental Figure 2). Overall, while there was near perfect separation of control samples from cancer samples using CelFiE-ISH with or without clipping (Figure 1C), the clipping version provided a more accurate representation of the true cell-type composition.

To evaluate the accuracy of tumor fraction estimation from cfDNA in CRC samples, we estimated tumor fractions from copy number analysis using ichorCNA (Figure 1D), and compared these to epithelial fractions estimated by methylation-based deconvolution (Figure 1E). CelFiE-ISH with clipping achieved lower root mean square error than other methods (RMSE = 0.099). This result was significantly better than the default CelFiE-ISH method (RMSE = 0.126), a difference that was largely driven by inflated epithelial fractions in samples with less than 10% tumor content (as seen in the “CelFiE-ISH” scatterplot in Figure 1E). CelFEER (RMSE=0.125), UXM (RMSE=0.195), and CelFiE (RMSE=0.104) also suffered from a degree of inflated epithelial fraction estimates. While CelFiE and CelFiE-ISH with clipping performed best for absolute RMSE (Figure 1F), the default CelFiE-ISH performed best for the correlation R^2^ (Figure 1G). Overall, all of the CelFiE-derived algorithms performed reasonably well.

### Pan-Epithelial Markers Improve Accuracy of cancer fraction estimation

Based on the improved performance of CelFiE-ISH with clipping, we concluded that many fragments could not be assigned unambiguously to a specific epithelial cell type. However, given that different epithelial cells have a large number of “pan-epithelial” methylation markers in common (3), we reasoned that using larger numbers of these pan-epithelial markers could give improved epithelial cell fractions. While it would in fact be valuable to distinguish between these different cell types, it may be too challenging given the very low genomic coverage here. This generalized “collapsing” approach may be useful for other cell lineages as well, as has been proposed previously (7).

We re-ran marker selection by collapsing each cell type group (lymphocyte, monocyte/macrophage, EP/MK, granulocyte, endothelial, hepatocyte, and epithelial) and picking markers for that group. We then performed deconvolution, at first using only the top 250 markers per cell type for consistency with our earlier analysis. The first thing we noticed was that the over-inflation of epithelial cell estimates was almost completely abrogated for all deconvolution methods (Figure 2A-B). Separation between control and cancer was improved for every deconvolution method, with UXM, CelFEER, and CelFiE-ISH (with or without clipping) all with area under the ROC curve (AUROC) above 0.99 (Figure 2B). The collapsed epithelial fraction estimates were significantly closer to ichorCNA-based estimates, compared with our previous 31-cell-type estimates (Figure 2C). RMSE values had a median value of 0.082 vs. a 31-cell-type value of 0.126 (Figure 2D), while R² values had a median value of 0.810 vs. a 31-cell-type value of 0.684 (Figure 2E). CelFiE-ISH (with or without clipping) and CelFEER had the best RMSE scores (∼0.08), whereas CelFiE and UXM performed more poorly (RMSE of 0.09 and 0.13, respectively). CelFiE-ISH performed only slightly better with clipping (RMSE of 0.077 vs. 0.082 without clipping).

**Figure 2.**
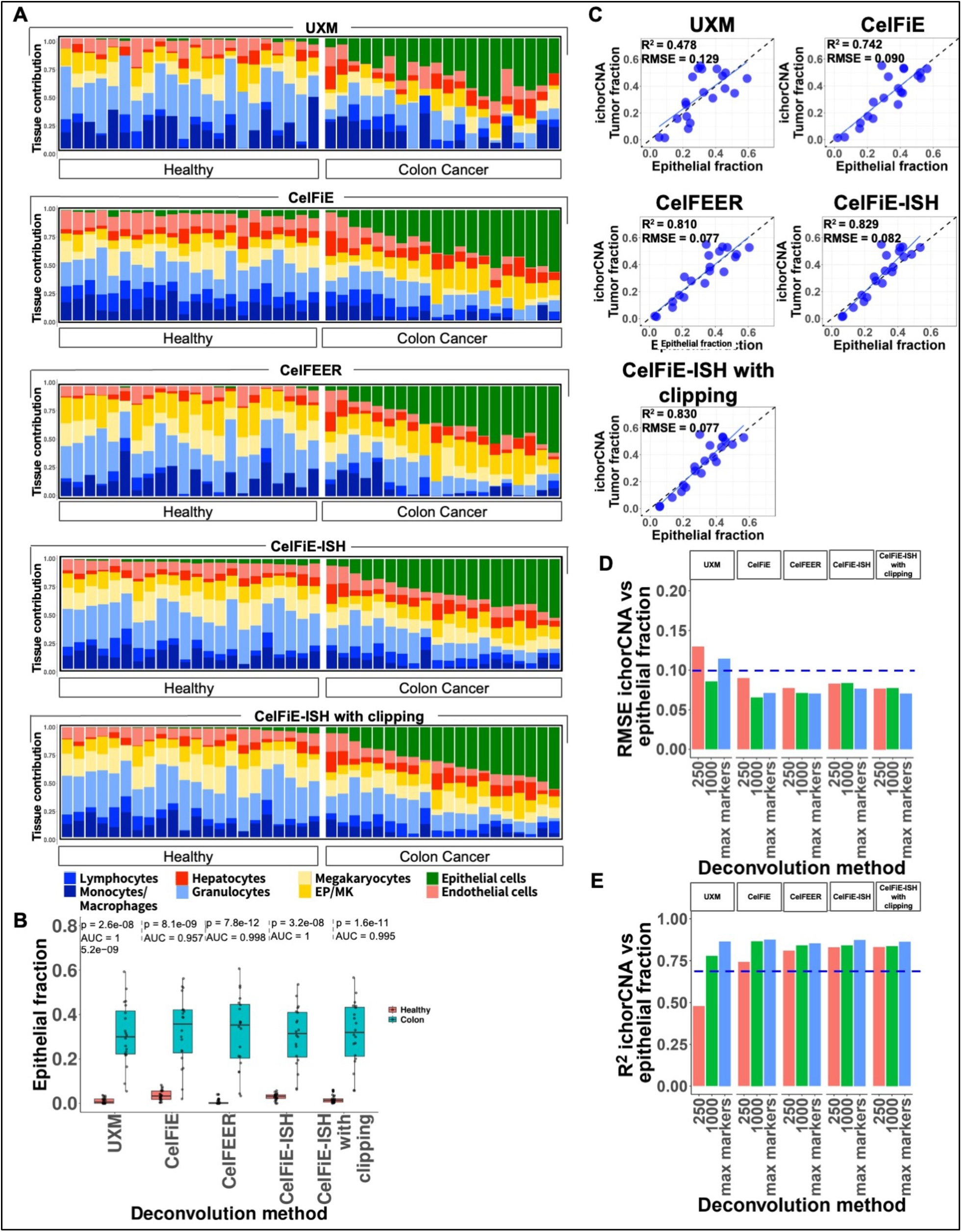
Pan-Epithelial Markers Improve Accuracy of cancer fraction estimation. **(A)** Cell-of-origin contributions from deconvolution using 250 markers from each of seven cell type groups commonly found in blood cfDNA (lymphocytes, monocytes, erythroid progenitors, granulocytes, endothelial cells, hepatocytes) plus a single pan-epithelial cell type group. Samples follow the same order as Figure 1. **(B)** ROC analysis comparing epithelial fractions between healthy and cancer samples for each deconvolution method. AUROC and *p*-values (one-tailed Wilcoxon test) are indicated. **(C)** Comparison of the collapsed epithelial fraction from each deconvolution method against ichorCNA tumor fractions. **(D–E)** Performance metrics across differing marker counts per cell type (250, 1,000, and “max markers”) for each deconvolution method, showing **(D)** RMSE and **(E)** R^2^.

To investigate the impact of marker set size, we repeated our analysis using collapsed marker sets but progressively increased the number of markers selected. Specifically, two distinct marker sets were employed: one comprising 1,000 markers per cell type and another containing the maximum possible 2,825 markers per cell type (see Methods). The latter set, hereafter referred to as the “max markers” set, was designed to ensure the highest possible and equal representation of markers across all cell types (see Methods). Of the methods that performed reasonably well under the 250 marker analysis (CelFiE, CelFEER, and CelFiE-ISH), adding additional markers reduced RMSE by 20% (Figure 2D), with similar minor improvements in R² (Figure 2E). We conclude that for most of the relatively high tumor fraction samples in this dataset, 250 markers per cell type perform relatively well for accurate deconvolution.

### The accuracy and limit of detection of epithelial fraction estimates with low tumor fraction

The analyses described above were performed using cfDNA extracted from plasma of mostly advanced colorectal cancer patients, which often release large quantities of cfDNA into circulation. In order to simulate lower tumor fractions (typical of many early stage cancers), we diluted each CRC sample *in silico* with reads from pooled healthy samples (Figure 3A). This dilution produced five additional versions of each sample in addition to the original (undiluted) sample, each with a cancer DNA fraction half of the previous dilution. In total, we produced 5*20 or 100 virtual samples in addition to the original 20 samples.

**Figure 3:**
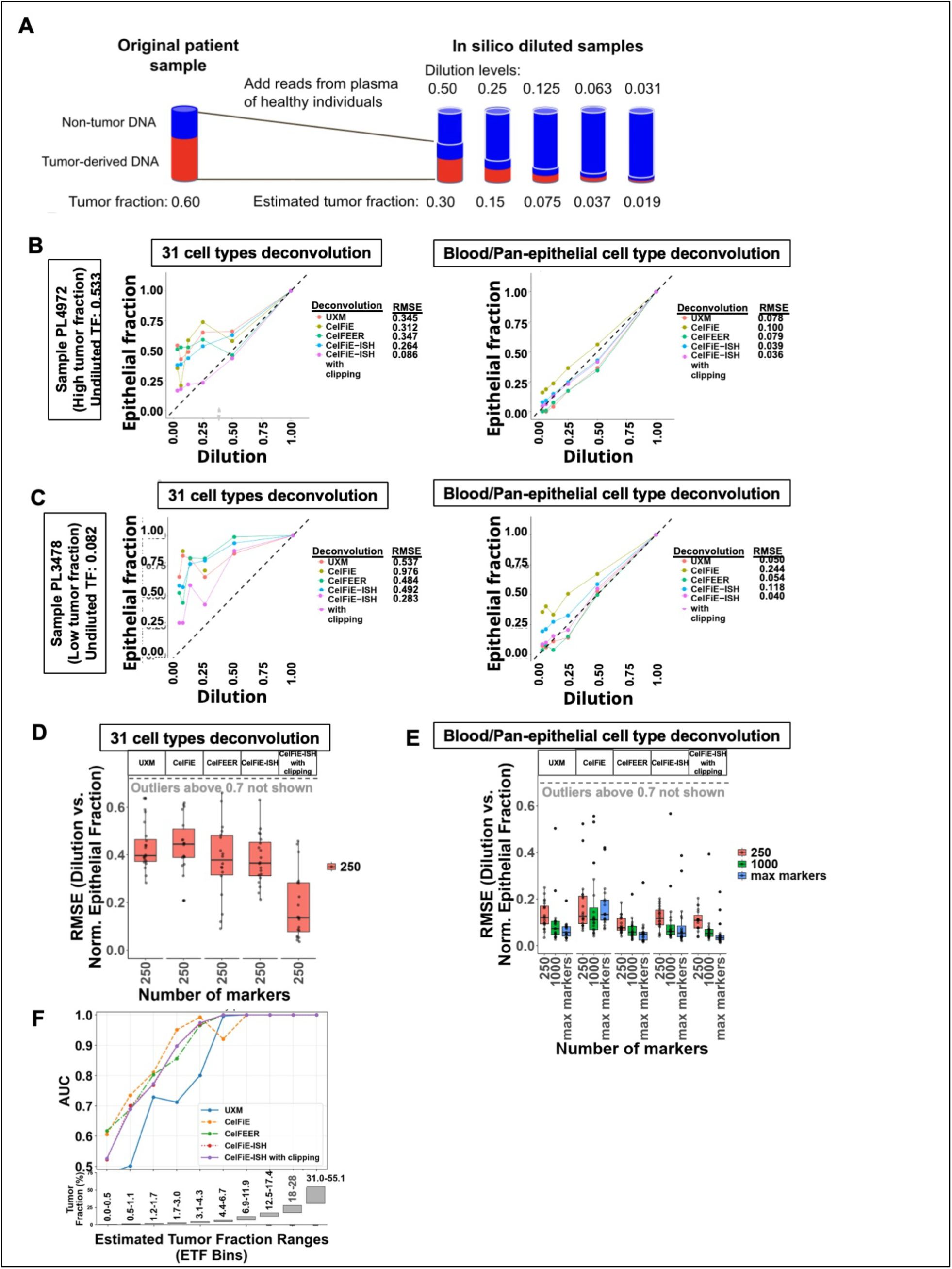
The accuracy and limit of detection of epithelial fraction estimates with low tumor fraction. **(A)** Schematic of the *in silico* dilution workflow. Twenty colorectal cancer samples were diluted with pooled healthy plasma using a two-fold serial dilution series to generate mixtures with reduced tumor content (ranging from 1.0 to 0.031 of the original cancer fraction). **(B)** Normalized epithelial fractions for a high tumor fraction sample (original TF = 0.533). The x-axis shows the estimated tumor fraction (original tumor fraction * dilution level) and the y-axis shows the normalized epithelial fraction (scaled such that the undiluted sample equals 1.0). The diagonal line represents the expected two-fold decrease. Comparisons are shown for the 31-cell-type marker set (left) and the pan-epithelial marker set (right). **(C)** Dilution analysis as in (B) for a sample with a lower tumor fraction (original TF = 0.082). **(D–E)** RMSE values calculated from the expected diagonal for all 20 colorectal cancer samples using either **(D)** the 31-cell-type marker set or **(E)** the pan-epithelial marker set with varying marker counts. **(F)** AUROC values for cancer vs. healthy classification across ten tumor fraction bins. Each bin contains 12 virtual colorectal cancer samples compared against 22 healthy controls (detailed analysis in Supplemental Figure 5). The box plot below the x-axis indicates the range of tumor fractions within each bin.

Since methylation-based deconvolution was highly accurate for the undiluted samples (Figure 2C), we reasoned that these were reliable and could be used to evaluate the relative tumor fractions of diluted samples. We assigned the undiluted sample an epithelial fraction value of 1, and plotted the relative epithelial fractions of all diluted samples (for a single “high tumor fraction” example deconvoluted using 31 cell types, as shown in Figure 3B). Since we expect each dilution to result in a two-fold decrease, the normalized epithelial fraction is expected to follow the diagonal line, but we observed significant overestimation for all deconvolution methods except CelFiE-ISH with clipping (Figure 3B). In contrast, the collapsed cell type deconvolutions performed quite well (Figure 3B, right). We can quantify this by calculating the RMSE vs. the expected diagonal line, which yielded a median RMSE=0.312 for the 31-cell-type deconvolutions vs. a median RMSE=0.078 for the collapsed cell type deconvolutions.

While the “high tumor fraction” sample above had a minimum diluted tumor fraction of 0.017 (original tumor fraction of 0.53 times 3.125%), we can investigate even lower tumor fractions with dilutions of samples that begin with lower tumor fraction in the original sample. As an example, we show a sample with an undiluted tumor fraction of 0.082 (Figure 3C). Here, we can see that even in the collapsed cell type deconvolution (Figure 3C, right) some of the estimates are quite poor at the middle 0.125 dilution level (which corresponds to a tumor fraction of 0.01 for this sample).

With many of the samples having diluted tumor fractions of 0.01 or less, we had an opportunity to investigate how well different deconvolution variants perform at low tumor fraction, using the RMSE from the diagonal line as a metric. Interestingly, CelFiE-ISH with clipping was the only deconvolution method that performed relatively well using the default 31-cell-type markers (Figure 3D and Supplemental Figure 3), underscoring the value of clipping in this context. Using the collapsed cell type markers, most of the deconvolution methods performed much better (Figure 3E and Supplemental Figure 4). As the number of markers increased, RMSE values decreased, particularly with 1,000 markers or more, highlighting the benefit of larger marker sets in the context of low tumor fraction and ultra-low genomic coverage. Overall, CelFEER and CelFiE-ISH (with or without clipping) performed markedly better than the other two methods. Accuracy was always somewhat higher in the max markers condition than the 1000-marker condition, where CelFiE-ISH with clipping achieved the absolute lowest RMSE of 0.034.

We also used this diluted sample set to estimate the limit of detection (LoD) in cancer vs. healthy comparisons. We binned each of the 120 diluted and undiluted samples into ten equally-sized bins based on their estimated tumor fraction (dilution level * undiluted tumor fraction). We then performed deconvolution using collapsed marker sets, and performed an ROC analysis of epithelial fraction in the samples in each bin vs. the 22 undiluted non-cancer control samples (Supplemental Figure 5 and Figure 3F). For CelFEER and CelFiE-ISH (with or without clipping), samples with tumor fractions of 1.7-3.0% had good classification (AUROC=0.85-0.95), whereas samples with tumor fractions 3.1-4.3% had close to perfect separation (AUROC=0.97-0.99). All had >0.99 separation for bins above 4.3%. CelFiE had somewhat inconsistent results, with better AUROC scores with samples lower than 4.3%, but poorer AUROC scores for samples >4.3%. UXM had poorer AUROC than other methods under all conditions. The ability to discriminate almost all cancer samples from non-cancer controls at a level of 3.1-4.3% tumor fraction is similar to the generally-accepted LoD of ichorCNA, which is ∼3%. (11, 13). Having two reliable but independent features for measuring tumor fraction from the same sequencing assay is a benefit of ONT WGS, since each feature alone is subject to error and noise.

### Using epithelial fraction to detect cancer DNA in other epithelial cancer types

In our colorectal cancer benchmarking, collapsed cell-type markers using the maximum number of markers using CelFiE-ISH with clipping outperformed other deconvolution methods. Thus, we used these settings to analyze a set of pre-treatment samples from several other epithelial cancer types (Supplemental Table 1). We compared our 22 non-cancer control samples to these breast cancer (n = 66), lung cancer (n = 6), and pancreatic cancer (n = 10) samples (Figure 4A). For the breast cancer samples, we calculated control vs. cancer AUROC values for each stage independently (Figure 4B). As with earlier reports (10, 14), stage IV cancers were readily detectable whereas earlier stages were not. The lung and pancreatic cancer cohorts were primarily stage IV cancers and performed well vs. non-cancer controls, with stage IV AUROC values of 0.836 (p = 0.019) and 0.916 (p = 0.00042), respectively (Figures 4C-D).

**Figure 4.**
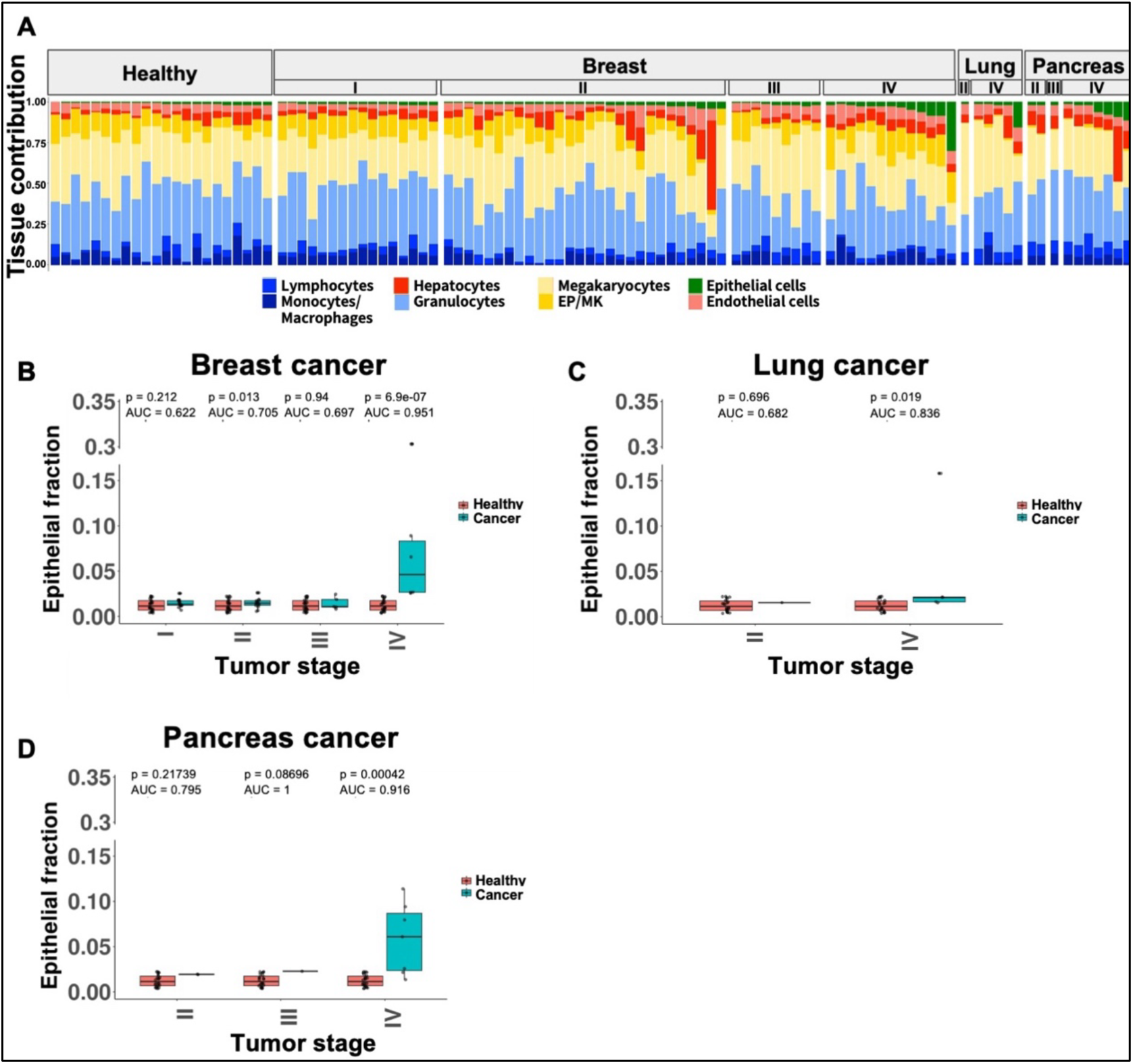
Using epithelial fraction to detect cancer DNA in other epithelial cancer types. **(A)** Cell-of-origin contributions estimated using CelFiE-ISH with clipping (pan-epithelial marker set, “max markers”) for healthy (n = 22), breast cancer (n = 66), lung cancer (n = 6), and pancreatic cancer (n = 10) cohorts. Samples are stratified by tumor stage and sorted by epithelial fraction (sample identifiers are provided in Supplemental Figure 6). **(B–D)** Comparison of epithelial fractions between healthy and cancer samples for **(B)** breast cancer, **(C)** lung cancer, and **(D)** pancreatic cancer. AUROC and *p*-values (one-tailed Wilcoxon test) were calculated for each cancer stage independently.

## Discussion

In this study, we demonstrated that ultra-low-pass whole-genome methylation sequencing using ONT can achieve a circulating tumor DNA (ctDNA) limit of detection (LOD) of approximately 1.7-3.1%. While this threshold may not support use for early detection, it provides an analytical performance comparable to ichorCNA, the current benchmark for copy-number alteration (CNA) quantification. This epigenetic approach is particularly valuable for malignancies characterized by few or no somatic CNAs, as such cancers remain largely undetectable using CNA analysis alone. Furthermore, profiling the methylome is critical for monitoring tumor evolution and detecting molecular switches that may suggest changes in treatment. Molecular subtype switching is a frequent driver of therapeutic resistance, exemplified by the conversion of estrogen receptor (ER)-positive to ER-negative disease in breast cancer (∼20–25%) (15), the transformation of non-small cell lung cancer (NSCLC) to small cell lung cancer (SCLC) (3–14%) (16), and the transdifferentiation of androgen receptor-active prostate cancer (ARPC) to neuroendocrine prostate cancer (NEPC) (15–20%) (17).

We addressed an inherent limitation of our Expectation-Maximization model noted during the earlier: the tendency to over-inflate estimates for rare or biologically absent cell types when represented by low-information reads. By implementing a probabilistic "clipping" strategy, we reduced spurious epithelial "noise" in healthy individuals to near-baseline levels and brought tumor fraction estimates in cancer samples into closer alignment with ichorCNA benchmarks. While this modification did not markedly shift the limit of detection for advanced colon cancer cases, it yielded a more faithful representation of global cell-of-origin compositions across diverse clinical cohorts. Consequently, we have integrated this clipping functionality into the public CelFiE-ISH package to improve general deconvolution accuracy.

We subsequently explored strategies to enhance the assay’s limit of detection (LOD), which is inherently constrained by read sparsity in ULP-WGS. Our default CelFiE-ISH package adopts a granular approach, treating each epithelial tissue independently and assigning 250 specific markers to each, which may be insufficient for ULP-WGS. To address this, we evaluated the use of "pan-epithelial" markers—genomic regions that consistently reflect epithelial lineage identity across diverse carcinomas without necessarily differentiating between specific epithelial cell types. While such markers might not support the definitive classification required for multi-cancer early detection (MCED) in a single step, they provide superior sensitivity for initial screening, which can be followed by a targeted analysis of markers that differentiate cancer types. This hierarchical strategy could be extended to other lineages, utilizing mesenchymal markers for sarcomas or neural signatures for central nervous system malignancies.

Indeed, adopting a lineage-specific detection framework yielded significant gains in both sensitivity and accuracy. This improvement was most pronounced in our *in silico* dilution experiments designed to simulate low-tumor-fraction scenarios. In these contexts, expanding the feature space to over 1,000 pan-epithelial markers proved pivotal, substantially reducing error rates and lowering the analytical limit of detection. We achieved robust separation between cancer and healthy samples at tumor fractions exceeding 1.7% (AUROC ∼0.9), with near-perfect discrimination at fractions above 3.1%. Given our ultra-low-pass sequencing depth (∼0.25x coverage), these results matched or exceeded the performance of established CNA-based pipelines like ichorCNA, which typically report a 3% detection floor (11, 13). As we have previously proposed (12), the integration of methylation and copy-number features from the same shallow WGS run could further refine detection thresholds and clinical utility.

## Conclusions

Our findings underscore that marker selection remains the most critical—and still evolving—aspect of methylation-based diagnostic inference. Genomic methylation markers span a range of specificities from highly cell-type-specific to restricted to either broad developmental lineages or ad hoc combinations of cell types (7). The effective selection of markers is determined by the interplay between biological specificity, sequencing depth, and physical read length. For instance, ∼167 bp fragments characteristic of cfDNA will require a larger number of independent markers than genomic tumor DNA sequenced with long-read sequencing such as ONT (15-20 kb). In this case, a smaller number of markers that span longer genomic intervals (such as super-enhancers) might be sufficient to provide highly sensitive and specific detection. We envision future strategies where different marker panels are pre-selected based on read lengths and genome coverage, or bespoke marker sets are determined on-the-fly for each dataset. Incorporating copy number into 5mC deconvolution strategies, or 5-hydroxymethylation (18, 19) (readily detectable via Oxford Nanopore sequencing) can help push the detection limit of ultra-low-pass WGS sequencing even further.

## Code availability

Most analysis was performed using publicly available tools, with all software version numbers and command line settings listed in the Methods section. For several in-house scripts that were developed to analyze fragmentation patterns, source code is deposited at Zenodo with DOI 10.5281/zenodo.18100426. This record is under embargo while this manuscript undergoes confidential peer review.

## Data availability

To enable full reproducibility of all analyses while preventing the disclosure of personally identifying genetic variants, two complementary data types were deposited. First, BAM files generated prior to filtering were anonymized by replacing all single-nucleotide variants with the corresponding hg38 reference bases and all insertion variants with Ns (see Methods). These anonymized BAM files preserve alignment structure and support all downstream filtering and copy number analyses without exposing individual-specific genetic information. These data are available in the Short Read Archive under BioProject ID PRJNA1394862.

Second, to provide read-level DNA methylation information, Biscuit (20) was used to generate epiBED files (see Methods). epiBED files represent each read on a separate line and encode all DNA methylation calls along the read, enabling integrated methylation analyses. These files are compatible with the CelFiE-ISH framework (6), which was used for cell-of-origin deconvolution. epiBED files are deposited in Zenodo (DOI: 10.5281/zenodo.18100426).

The CelFiE-ISH deconvolution model with clipping support is available on GitHub (https://github.com/methylgrammarlab/deconvolution_models), and all input files required for deconvolution using blood and pan-epithelial cell types are provided in the associated Zenodo repository (DOI: 10.5281/zenodo.18100426).

## Author contributions

AZ, BPB, and AE conceived the study. AG, AZ, BU, TP, LK, AH, JC, AS, MT and TS enrolled patients, did clinical data entry, and performed clinical procedures. AG (Albert Grinshpun) and AZ coordinated sample collection and processing, with MM, AG (Ahinoam Gugenheim) and NS collecting and processing blood samples, and MI, MTM, SAK, and EK assisting with clinical database construction and analysis. SI performed ONT sequencing and AD performed base-calling and alignment. EK performed all downstream computational analysis and produced final paper figures with BPB and AE supervising. IU assisted with changes to the CelFiE-ISH package. EK and BPB wrote the manuscript and AE assisted. BPB and AE supervised all aspects of the project.

## Acknowledgements

We thank Judith Magenheim, Yuval Dor, Ilana Fox-Fisher, and Ruth Shemer for providing non-cancer controls. We thank Netanel Loyfer for assistance with wgbstools and DNA methylation atlas datasets. Computation was performed on the Hebrew University Research Computing Services cluster.

## Ethics approval and consent to participate

The study was approved by the Ethics Committees of the Hadassah-Hebrew University Medical Center of Jerusalem #0346–12 allowing identification of individual participants during or after data collection. Informed consent was obtained from all individuals before blood sampling and complied with the Declaration of Helsinki and the principles of Good Clinical Practice guidelines.

## Methods

### Patient Cohort

The study population consisted of either patients diagnosed with colon, pancreatic or lung cancer, before oncological treatment or women with a breast mass referred for a breast biopsy, before the biopsy was performed. All patients were treated at the Hadassah Medical Center. Sample information is shown in Supplemental Table 1, and subject/patient information is shown in Supplemental Tables 2-6.

### Sample preparation and DNA extraction

Peripheral blood samples were collected in Streck tubes and centrifuged for 10 min (890 × g at 4°C for EDTA-containing tubes or 1400 × g at 23°C for Streck). The supernatant containing plasma was transferred to Eppendorf tubes without disturbance of the cellular layer and centrifuged at 16000 × g for 20 min at 4°C. The supernatant then was collected in 2-ml Maxymum recovery tubes (Axygen) and stored at −80°C. cfDNA was extracted from 2 to 4 ml of plasma using the cfDNA Serum and Plasma Kit (Zymo Research) or a QIAsymphony liquid handling robot (Qiagen), as described in (21). DNA quantity was estimated using the Qubit dsDNA HS assay kit (Thermo Fisher Scientific).

### Library preparation and Sequencing

The original sample names, barcode and kit used from that study are listed in Supplemental Table 1. Library preparation was done based on ONT protocol with minor changes. We used 3ng input cfDNA from each sample.

End repair and A-tailing used NEBNext^®^ FFPE DNA Repair Mix (NEB #M6630) and NEBNext^®^ Ultra™ II End Repair/dA-Tailing Module (NEB #E7546), according to NEB protocol: 12 uL of cfDNA, 0.875 uL of NEBNext FFPE DNA Repair Buffer, 0.875 uL of Ultra II End-prep reaction buffer, 0.5 uL NEBNext FFPE DNA Repair Mix were incubated in a thermocycler at 20°C for 30 minutes (with the heated lid off). Next, 0.75 uL of NEBNext Ultra II End Prep enzyme mix was added and placed in a thermocycler (with the heated lid set to 75°C) with the following program: 30 minutes at 20°C, 10 minutes at 65°C, and hold at 4°C. DNA was cleaned with 2.3x magnetic beads (Agencourt AMPure XP, BC-A63881), washed 2x with 80% ethanol and eluted with 11ul Nuclease-free Water (NEB #B1500).

Barcode ligation used Blunt/TA Ligase Master Mix (NEB #M0367) with 1.5 uL of Native Barcode in total volume of 25 uL. DNA was cleaned separately with 0.8x magnetic beads (Agencourt AMPure XP, BC-A63881), washed 2x with 80% ethanol and eluted with 11 uL Nuclease-free Water (NEB #B1500). Samples were pooled and Native Adapter was added according to ONT protocol, with the SFB (short fragment buffer).

### High accuracy calling (HAC) base calling and alignment of Nanopore fast5 files

All CRC and breast cancer Fast5 samples were base called and demultiplexed using Guppy (version 6.3.8+d9e0f64) with the dna_r10.4.1_e8.2_400bps_modbases_5mc_cg_sup.cfg model and parameters including “--barcode_kits SQK-NBD114-24 --trim_adapters --do_read_splitting.” Alignment for these samples was performed using minimap2 (version 2.22-r1101) with default parameters and “map-ont” set to ONT. Lung and pancreas samples were processed entirely with Dorado (version 0.7.2+9ac85c6), which handled both base calling and alignment using its internal minimap2-based aligner (DS:2.27-r1193). Healthy and other non-cancer control samples were split evenly between the Guppy and Dorado pipelines. An overview of sample processing is provided in Supplemental Table 1. All resulting BAM files were used in downstream methylation analysis and quantification workflows. All alignments were performed relative to the hg38 genome build.

### Marker selection for 31 cell types

For the 31 cell type deconvolution, we used the cell-type specific regions identified by Loyfer et al., using the top 250 markers (the “U250” set) per cell type, that included a maximum of 250 markers per cell type.https://github.com/nloyfer/UXM_deconv/blob/main/supplemental/Markers.U250.hg38.tsv. This set included 40 cell types (9521 regions). We removed regions that fall within sex chromosomes and used marker regions from 31 cell types, as in (6), reducing the total number of marker regions to 7451. For visualization purposes, individual cell types were aggregated into eight broader groups. Monocytes/Macrophages included blood monocytes and tissue macrophages. Lymphocytes included B cells, T cells, and NK cells. The Erythroid Progenitor/Megakaryocyte (EP/MK) lineage was grouped together because the marker set lacked megakaryocyte-specific markers, and erythroblast-derived cfDNA signals in plasma are thought to predominantly reflect megakaryocyte origin (22). Additional groups included Neutrophils/Granulocytes; Hepatocytes; Epithelial cells (bladder, breast basal and luminal, colon, fallopian tube, stomach, head and neck, kidney, lung alveolar and bronchial, ovary, endometrium, prostate, small intestine, thyroid, pancreas acinar and ductal); Endothelial cells; and an “Other” category containing cardiomyocytes, neurons, oligodendrocytes, adipocytes, and pancreatic alpha, beta, and delta cells. Full collapsing of individual cell types into groups is shown in Supplemental Figure 2.

### Marker selection for blood vs. pan-epithelial cell types

Cell types were aggregated into eight collapsed groups as follows. The Blood-Granulocyte group comprised blood granulocytes (n=3). The Blood-Monocyte/Macrophage group included blood monocytes (n=3) and macrophages derived from colon (n=2), liver (n=1), lung alveolar tissue (n=2), and lung interstitial tissue (n=3). The Endothelial group included endothelial cells from aorta (n=1), saphenous vein (n=3), kidney glomerular tissue (n=3), kidney tubular tissue (n=3), liver (n=1), and lung alveolar tissue (n=3). The Epithelial group incorporated epithelial cell types from bladder (n=5), breast basal (n=4) and luminal (n=3) epithelium, colon (n=8), endometrium (n=3), esophagus (n=2), fallopian tube (n=3), stomach (n=11), kidney (n=8), larynx (n=1), lung alveolar and bronchial epithelium (n=6), ovary (n=1), pharynx (n=1), prostate (n=4), small intestine (n=5), thyroid (n=3), tongue (n=4), and tonsil (n=5). Additional groups included Erythroid/Megakaryocyte (n=3), Hepatocytes (n=6), Lymphocytes consisting of blood B cells (n=5) and T cells (n=22), and Megakaryocytes/Platelets (n=3). Megakaryocyte/Platelet data is from (22).

For each cell type, whole-genome bisulfite sequencing (WGBS) BAM files were converted to PAT format using wgbstools suite (commit 3324a60e912cc36387c5f0b9f40ecc88a9678ef5, nloyfer/wgbs_tools) with the command wgbstools bam2pat -q 30 --lbeta --genome hg38. The resulting beta files were used for DNA methylation marker discovery using the wgbstools suite. Genomic methylation segments were generated with wgbstools segment --min_cpg 4 --pcount 30 using the beta files from all reference cell types. The resulting blocks file served as input for wgbstools find_markers to identify cell-type–specific methylation markers. Marker identification incorporated the beta files, the blocks file, and a predefined cell-type group file (available on GitHub) using the following parameters:

> --min_bp 0 --max_bp 10000000000 --min_cpg 0 --max_cpg 10000000000 --min_cov 4 --na_rate_tg 0.334 --na_rate_bg 0.334 --only_hypo --delta_means 0.001 --delta_quants 0.0 --tg_quant 0.33 --bg_quant 0.05 --unmeth_quant_thresh 0.25 --meth_quant_thresh 0.05 --unmeth_mean_thresh 0.25 --meth_mean_thresh 0.05 --top 10000 --pval 0.05. The --top 10000 setting was selected to maximize the number of candidate markers available for downstream filtering.

After marker identification, the resulting marker lists were combined, sex chromosomes and duplicate markers were removed (retaining for each duplicate the marker with the highest delta_quants and tg_mean values). To standardize the number of markers across all cell types, we reduced the maximum marker list for each cell type to match the cell type with the lowest initial number of markers (2,825 markers), thereby ensuring equal number of markers for “max markers” downstream analyses. From this set, we took the top 1,000 markers per cell type, or 250 markers per cell type, first by “delta quant” then by “tg_mean”.

### Deconvolution

For the deconvolution, we used the bam files after filtering with the same filtration as in (12): Samtools (Version 1.16.1) was used to filter BAM alignments, filtering out reads in blacklist regions (from: https://github.com/Boyle-Lab/Blacklist/blob/master/lists/hg38-blacklist.v2.bed.gz), unmapped reads, secondary and supplementary reads, reads with mapping quality less than 50, and reads longer than 700 bp. The later filtration was done to avoid including any DNA released during blood processing (23). For this, we used the pipeline in https://github.com/methylgrammarlab/cfnanoseq/tree/master.

For UXM deconvolution we used the code available at https://github.com/nloyfer/UXM_deconv (version 0.1.0). In order to convert the bam files to pat, we used wgbstools bam2pat (from wgbstools suite at https://github.com/nloyfer/wgbs_tools) with -F 0 -q 0 --nanopore. Then we build the atlas with uxm build, changing minimum CpGs 4 (--rlen 4). Then, we run deconvolution using uxm deconv, again requiring –rlen 4 as minimum CpGs in read to be considered.

To create the reference atlas for CelFiE and CelFiE-ISH, we used the pipeline in https://github.com/methylgrammarlab/cfnanoseq/tree/master

To get the bedgraph files needed to create the reference atlas, we first converted the cell types bam files from (3) to beta files with wgbstools bam2pat using “-q 30” to remove low quality reads. Then, pat files were converted to beta using “wgbstools pat2beta”. Then, beta files were converted to bed using “wgbstools beta2bed --genome hg38 --keep_na”. For each file, we kept only reads with full lines (5 fields), sorted, and reduced the end of the region by 1. These files were used as input to the pipeline, setting “--celfie_format_cell_types false –seperate_strands false”

For CelFiE deconvolution we used the code available (Version 1) at https://github.com/christacaggiano/celfie, with the following command:

~~~
celfie.py ${input} $output_dir 1 -u 0
~~~

For CelFiE-ISH deconvolution, we used the pipeline in (6) from https://github.com/methylgrammarlab/deconvolution_in_silico_pipeline, and https://github.com/methylgrammarlab/deconvolution_models v0.0.3.

For CelFiE-ISH with clipping, the --probs option was used to generate a probability matrix in which each read was assigned a probability of originating from each cell type. For clipping low-confidence cell type calls, we added the new flag --read_max_clipping_cell_type_probabilty and set it to 0.05, which clips all cell type probabilities less than 0.05 to 0. The command run was:

~~~
deconvolution --model celfie-ish -j {config} outfile={output} --probs
~~~

For CelFEER deconvolution, we implemented the CelFEER algorithm within the CelFiE-ISH framework (6). The primary modification involved calculating the methylation density of each fragment as the quotient of methylated cytosines relative to the total number of cytosines in the fragment. Following this calculation, fragment densities were discretized into five distinct bins (0-12.5%, 12.5–37.5%, 37.5–62.5%, 62.5–87.5%, and 87.5–100%) as required by the algorithm. These binned counts were subsequently aggregated into a sample-region matrix. Tissue-specific proportions were generated using CelFEER (https://github.com/pi-zz-a/CelFEER commit 58a1c60a30ae0161a8e2cb8624d73c035374cf1d) with the command:

~~~
python celfeer.py <input> <output_dir> 1.
~~~

### ichorCNA analysis for CRC samples

BAM files after filtering were used as an input. Somatic copy number analysis and tumor fraction estimates were performed using the ichorCNA package v.0.5.0 (11).

ichorCNA tumor fraction estimates are available in Supplemental Table 1. The ichorCNA parameters are, “normal = c(0.2,0.3,0.4,0.5,0.6,0.7,0.8,0.9,0.95,0.97,0.98,0.99), maxCN = 3, includeHOMD = False, estimateNormal = True, estimatePloidy = True, estimateScPrevalence = False, altFracThreshold = 0.05, flankLength = 100000, txnE = 0.9999999, chrTrain = ‘c(1:22)’, chrs = ‘c(1:22)’”.

We used a panel of normals (PoN) for ichorCNA analysis, using 42 healthy and “healthy-like” samples from our collection (specified by “Used in panel” column in Supplemental Table 1). These samples were chosen using an analysis of the wig files from ReadCounter (which are the input files for ichorCNA). We used ComplexHeatmap (24) for hierarchical clustering of the samples (shown in Supplemental Figure 1A). We found that nearly all of the healthy samples (20/22) clustered in a single cluster of 42 samples, which also included 22 early stage breast cancer samples, which are known to contain extremely low levels of ctDNA. We checked tumor fraction estimates of healthy samples with and without using this PoN, and found that tumor fraction only decreased (shown in Supplemental Figure 1B).

### Serial in-silico dilution of colorectal cancer samples

First, we generated a pooled healthy control sample by merging BAM files from all available healthy controls (n = 20) using the “samtools merge” command. From this pooled healthy sample, we randomly selected reads to complete a “non-cancer” component for each diluted sample as described below.

For each colorectal cancer (CRC) sample, a filtered BAM file representing the original, undiluted cancer sample (designated as 100% cancer content) was used as the primary input. For each dilution step, reads were randomly sampled from the original BAM file to produce a cancer component that was serially reduced by half −50%, 25%, 12.5%, 6.25%, and 3.125%. The remaining fraction of the reads for each dilution were randomly sampled from the pooled healthy sample, such that the total number of reads for the diluted sample was identical to the original undiluted cancer sample. The plot was generated using ggbreak (25).

### Estimating limit of detection relative to tumor fraction

For the limit of detection analysis, we used all undiluted and *in silico* diluted colorectal cancer samples. The estimated tumor fraction for each of these virtual samples was computed as the product of the original tumor fraction (estimated from ichorCNA on the undiluted sample) and its corresponding dilution percentage. Samples were then grouped into 10 bins based on estimated tumor fraction quantiles, and deconvolution was performed on each to determine the epithelial fraction. For each bin, we compared the epithelial fractions of the cancer samples in that bin to the full set of healthy controls (n=22) using the area under the receiver operating characteristic (AUROC) analysis.

### Anonymized BAM files

We used script bamReplaceSeqByRef.py with “-ref hg38.analysisSet.fa”, the same UCSC reference genome used for alignment. This script was adopted from (23). Files are available in the Short Read Archive BioProject ID PRJNA1394862, and code is available in the Zenodo repository listed in Code Availability.

### Creation of Biscuit EpiBED files

Filtered BAM files were used as input. Hydroxymethylated bases were converted to methylated bases using modkit (https://github.com/nanoporetech/modkit) v0.6.0 with the ‘modkit modbam adjust-mods’ command and the ‘--convert h m’ option. Hard clipping was removed from the start and end of CIGAR strings, and empty methylation tags were added to ensure compatibility with Biscuit. Epireads were generated using Biscuit v1.7.1 with the command “biscuit epiread -M -b 0 -m 0 -a 0 −5 0 −3 0 -y 0.9 -L 2300000 -E hg38.analysisSet.fa”, using the same UCSC reference genome as for alignment. Variant run-length–encoded strings were masked. All resulting files are available in the Zenodo repository described in the Data Availability section.

## Supplemental Figures

**Supplemental Figure 1.**
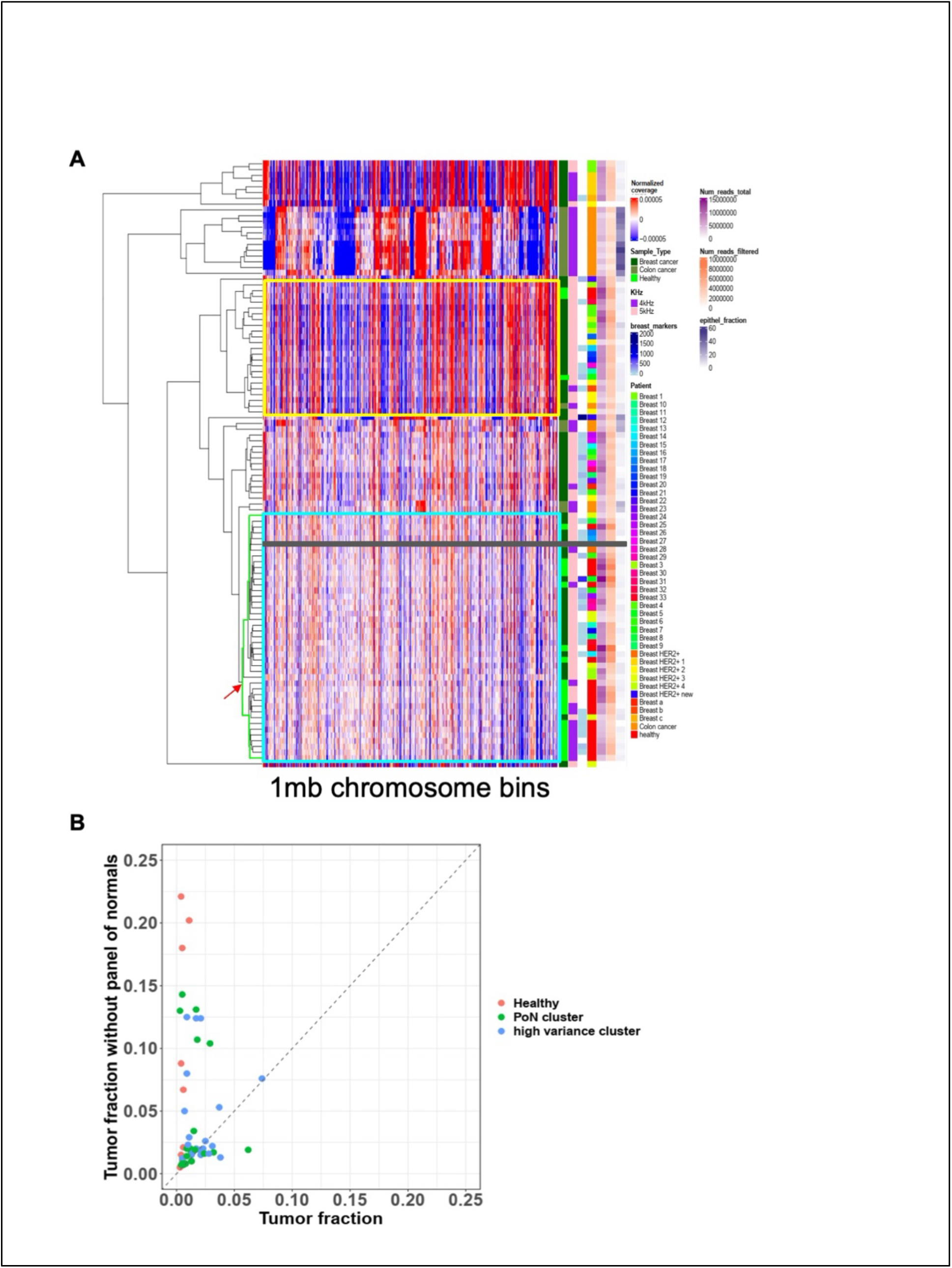
Panel of normals. **(A)** Hierarchical clustering heatmap of genome-wide coverage calculated in 1 Mb windows. Rows represent individual samples; columns correspond to genomic regions ordered by chromosomal position. Samples within the light blue rectangle were selected as normal references for ichorCNA; one colorectal cancer sample (grey rectangle) was excluded (see details in Methods section). The yellow rectangle marks samples with high variance (healthy samples within this cluster were not included). Sidebar tracks display normalized coverage, sample type, Nanopore read count, biological markers, patient ID, total and filtered read counts, and epithelial fraction (estimated using CelFiE-ISH with 250 markers using the pan-epithelial marker set). **(B)** Comparison of ichorCNA tumor fraction estimates with (**x-axis**) versus without (**y-axis**) using the reference panel of normals. Samples are colored by category, including breast samples and healthy controls that were included in the panel of the panel of normals and breast samples from the high-variance cluster (yellow rectangle in A).

**Supplemental figure 2.**
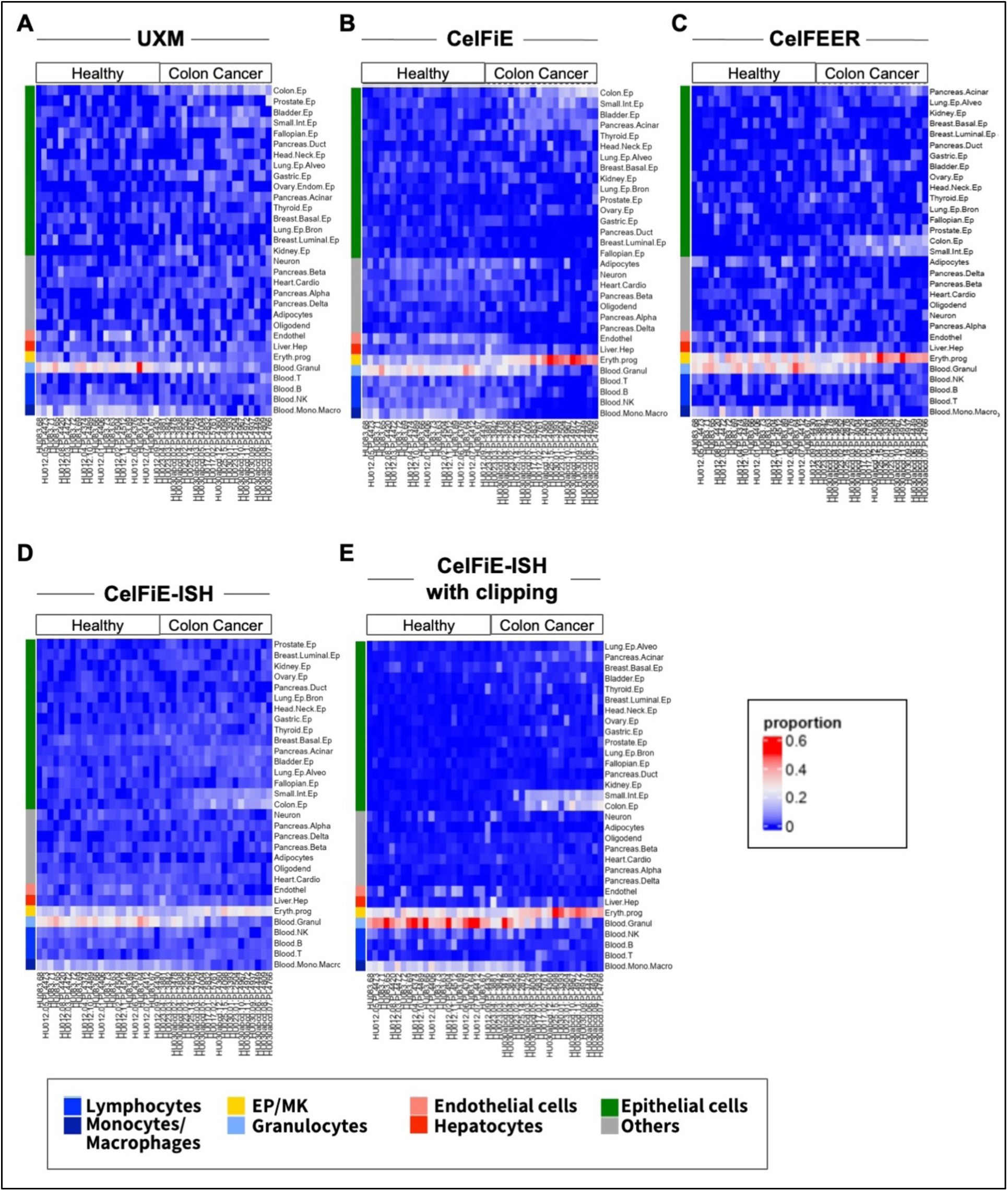
Cell type proportions from 31 cell-type marker set. Heatmap of cell-type deconvolution of 20 colorectal cancer samples and 22 healthy samples showing the 31 cell type marker set. All cell types are shown, and the cell type groups from Figure 1 are indicated.

**Supplemental figure 3.**
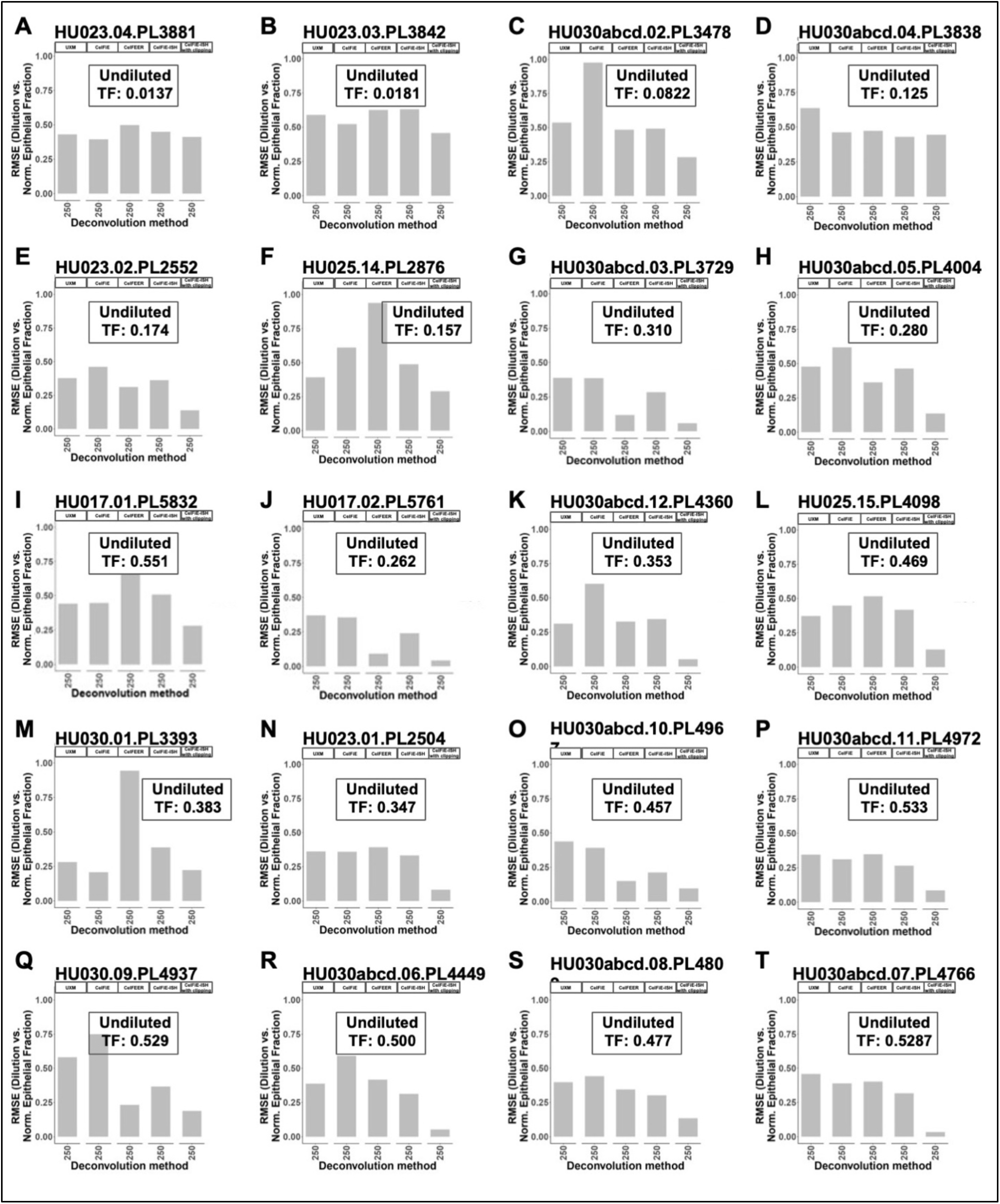
Dilution analysis error values, 31 cell-type marker set. Root mean square error (RMSE) for *in silico* diluted samples for each colorectal cancer case (n = 20), calculated using deconvolution with the 31-cell-type marker set. Samples are presented in the same order as in Figure 2A and Supplemental Figure 2.

**Supplemental figure 4.**
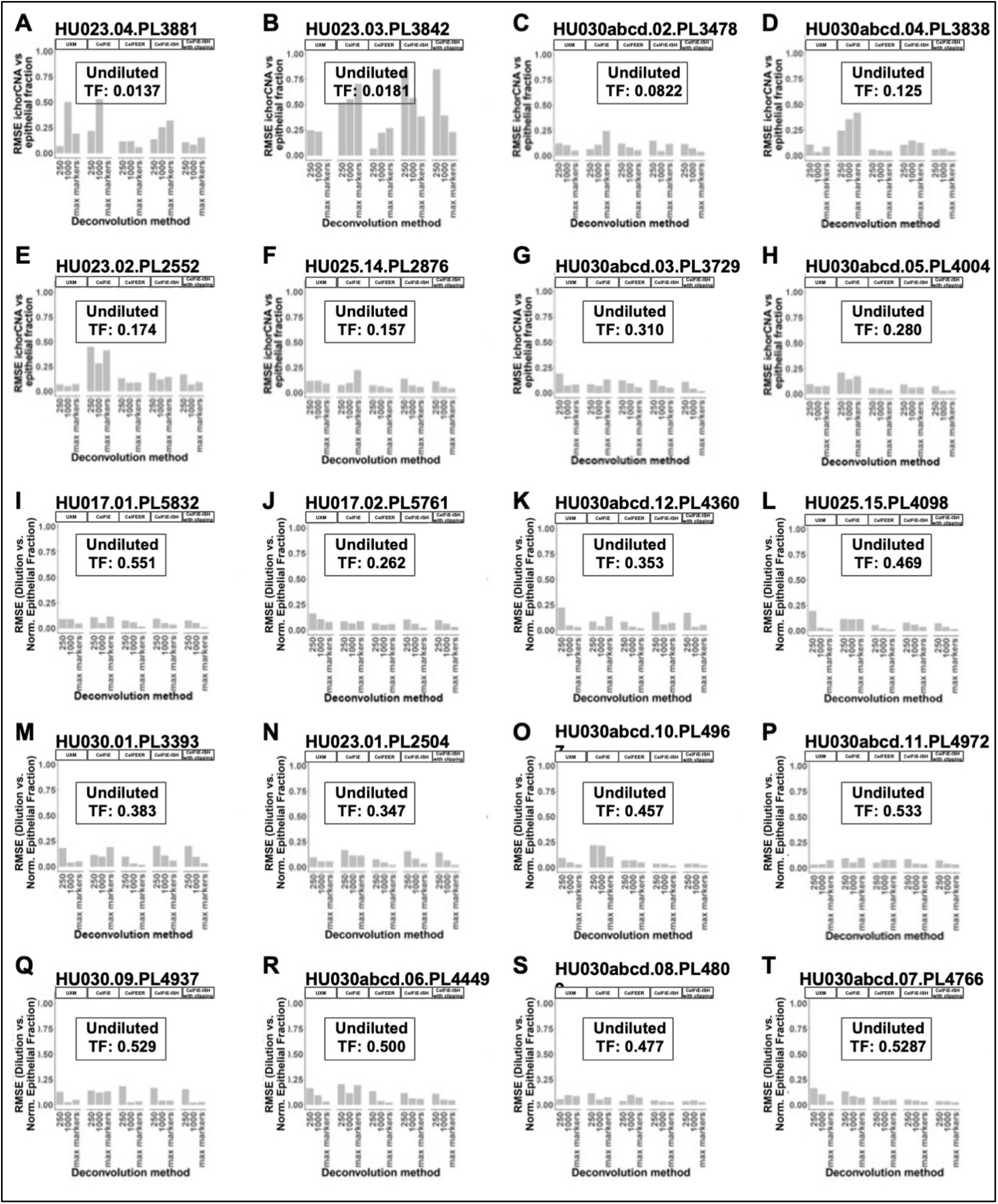
Dilution analysis error values, pan-epithelial marker set. Root mean square error (RMSE) for *in silico* diluted samples for each colorectal cancer case (n = 20), calculated using deconvolution with the pan-epithelial marker set. Samples are presented in the same order as in Figure 2A and Supplemental Figure 2.

**Supplemental figure 5.**
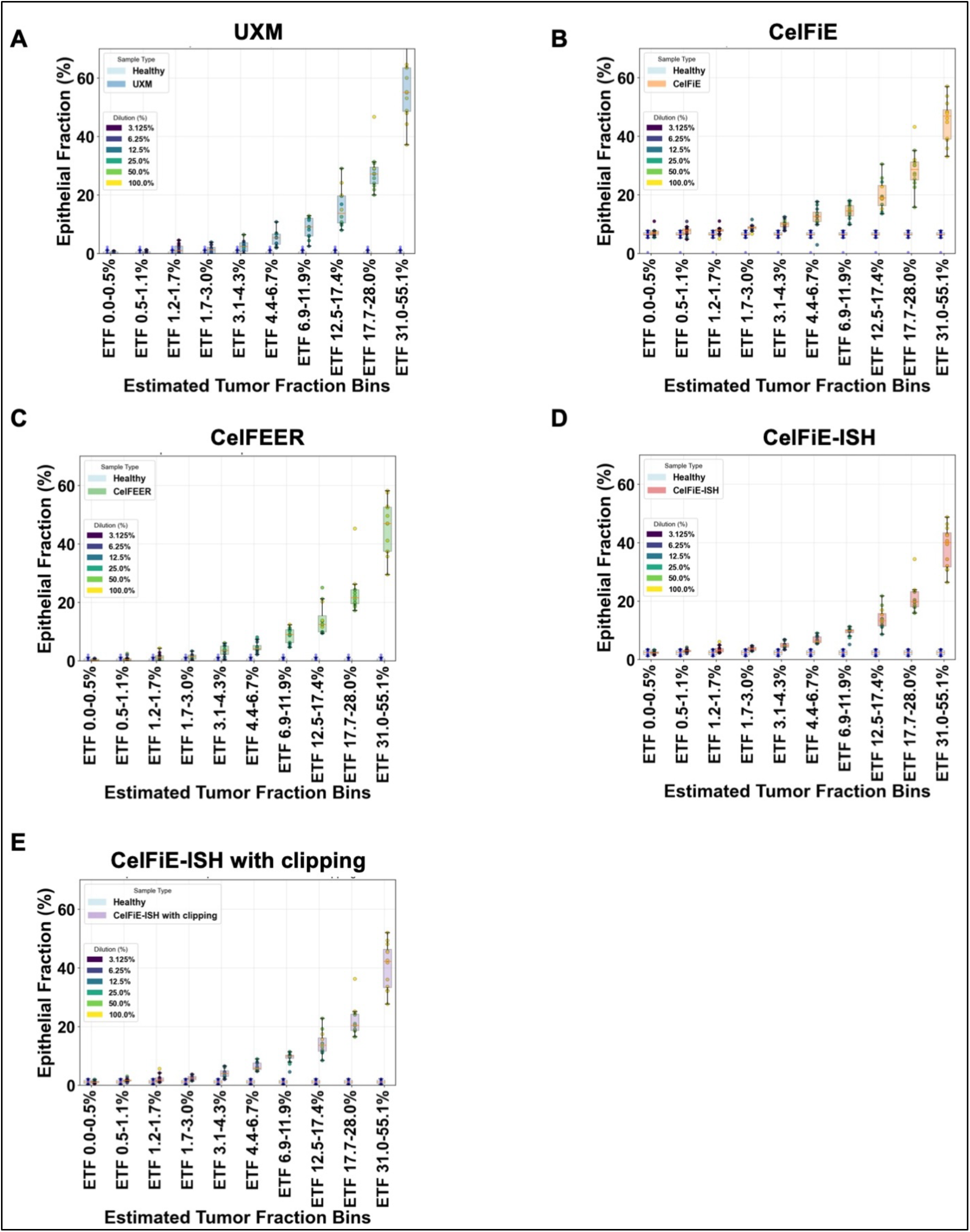
healthy vs. cancer comparisons stratified by tumor fraction. Each plot displays the 120 virtual colorectal cancer samples divided into 10 equal bins (n = 12 per bin), based on the estimated tumor fraction (ETF) values described in Figure 3F. For each bin, epithelial fractions estimated from deconvolution are shown for the 22 non-cancer control samples versus the 12 cancer samples within that bin. Individual samples are represented as points, colored by dilution level (where higher dilution corresponds to lower ETF). Each panel corresponds to a different deconvolution method, all using the pan-epithelial max marker set: **(A)** UXM, **(B)** CelFiE, **(C)** CelFEER, **(D)** CelFiE-ISH, and **(E)** CelFiE-ISH with clipping. These comparisons provide the underlying data for the AUROC analysis presented in Figure 3F.

**Supplemental figure 6:**
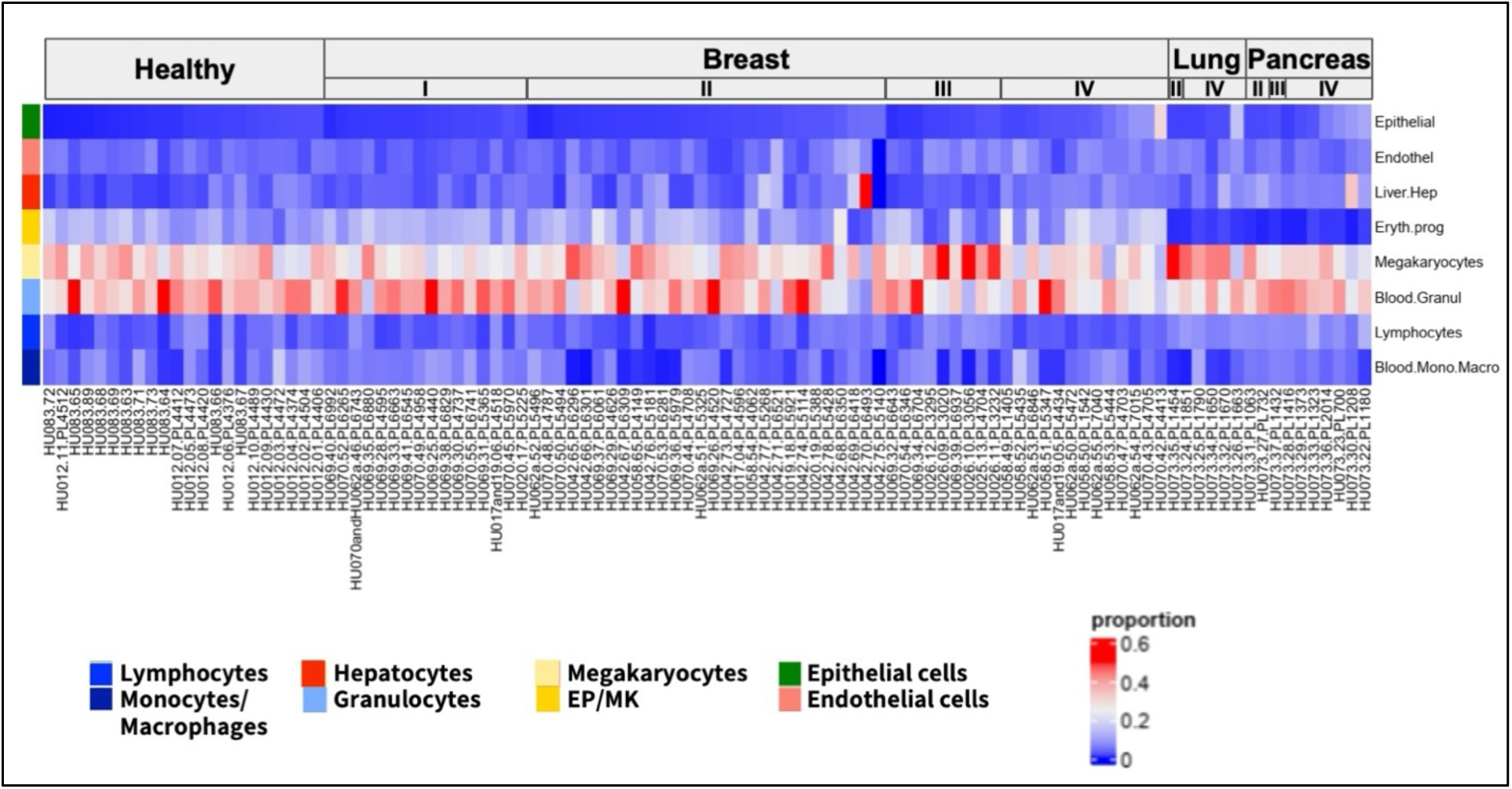
Sample ordering for Figure 4A. Same sample ordering as Figure 4A, showing sample IDs and deconvolution for all cell type groups for the pan-epithelia, max marker set.

## Supplemental Tables

Supplemental Table 1: Sequencing information and sequencing statistics for all samples included in the study.

Supplemental Table 2: Patient information for colorectal cancer patients included in study

Supplemental Table 3: Patient information for breast cancer patients included in study

Supplemental Table 4: Patient information for lung cancer patients included in study

Supplemental Table 5: Patient information for pancreatic cancer patients included in study

Supplemental Table 6: Patient information for non-cancer individuals included in study

Supplemental Table 7: Sequencing run information

